# Objective Classification of Neocortical Pyramidal Cells

**DOI:** 10.1101/349977

**Authors:** Lida Kanari, Srikanth Ramaswamy, Ying Shi, Sebastien Morand, Julie Meystre, Rodrigo Perin, Marwan Abdellah, Yun Wang, Kathryn Hess, Henry Markram

## Abstract

A consensus on the number of morphologically different types of pyramidal cells (PCs) in the neocortex has not yet been reached, despite over a century of anatomical studies. This is because of a lack of agreement on the subjective classifications of neuron types, which is based on expert analyses of neuronal morphologies: the shapes of somata, dendrites, and axons. Even for neurons that are visually different to non-experts, there is no common ground to consistently distinguish morphological types. We found that objective classification is possible with methods from algebraic topology, and that the dendritic arbor is sufficient for reliable identification of distinct types of PCs. We also provide a solution for the more challenging problem of whether two similar neurons belong to different types or to a continuum of the same type. Using this scheme, we objectively identify seventeen types of PCs in the rat somatosensory cortex. Our topological classification does not require expert input, is stable, and helps settle the long-standing debate on whether cell-types are discrete or continuous morphological variations of each other.

## Introduction

The mammalian neocortex is comprised of about 85% excitatory pyramidal cells (PCs), and around 15% inhibitory interneurons (Ramón y Cajal 1911; DeFelipe and Farinas 1992; Spruston 2007; Markram et al. 2015; Ramaswamy and Markram 2015). PCs, also termed principal cells, are characterized by a triangular soma, two distinct dendritic domains, both of which exhibit a high density of spines, emanating from the base (basal dendrites) and the apex of the soma (apical dendrites, respectively), and an axon that usually forms several local collaterals before leaving the neocortex to project to distant brain regions. Basal dendrites are localized around the soma while apical dendrites typically extend towards the pia, forming multiple oblique dendrites *en route* and terminating in a distinct tuft that is associated with high branching density.

Apical dendrites impart unique functional properties to PCs and form the basis for the generation of active dendritic (Cuntz et al. 2007, van Elburg & van Ooyen 2010, Cuntz 2012, van Ooyen & van Elburg 2014, Bird & Cuntz 2016) and synaptic events, such as back-propagating action potentials (Stuart & Sakmann, 1994), calcium transients in dendrites (Markram & Sakmann, 1994; Schiller et al. 1995; Yuste et al. 2000), integration of synaptic inputs from different cortical layers (Larkum et al. 1999; Larkum et al. 2001; Schaefer et al. 2003; Spruston, 2008), and spike-timing dependent plasticity (Markram et al. 1997a; Sjöström et al. 2001; Froemke et al. 2005). The unique functional properties of apical dendrites are therefore essential for integrating top-down (from association areas) and bottom-up streams of input (from primary sensory and motor areas) to the neocortex to shape the output firing pattern of PCs.

The characteristic morphological shapes of apical dendrites are associated with their unique functional properties, as objectively defined types of PCs also express unique firing patterns (Deitcher et al. 2017) and form distinct synaptic sub-networks within and across layers (Yoshimura et al. 2005; Kampa et al. 2006). Therefore, the branching properties of the apical trees are commonly used for their separation into morphological cell types (Ascoli & Krichmar 2000, Oberlaender et al. 2011, Marx and Feldmeyer 2012, Narayanan et al. 2017). The *expert classification*, which is based on visual inspection of the cells, usually makes it possible to distinguish the different shapes of morphologies and to group neurons into cell types. However, despite the expertise involved, visual inspection is subjective and often results in non-consensual and ambiguous classifications (DeFelipe et al. 2013, Ledergerber and Larkum 2010, Marx and Feldmeyer 2012, Markram et al. 2015). A striking indication of this problem, as described previously (DeFelipe et al. 2013), is the fact that experts assign a different cell type to a neuron from the one they had chosen in their original study for the same neuron, independently of the reconstruction quality (DeFelipe et al. 2013).

For this reason, an *objective classification* scheme is essential for a consensual and consistent definition of neuronal types, which can be achieved either by an *objective supervised* or *unsupervised classification* scheme. The *objective supervised classification* starts from the expert classification and verifies or disproves a proposed grouping based on objective measurements. When the expert classification cannot be supported by objective measurements, an *objective unsupervised classification* scheme is required. In this case, the classifier starts from a random classification and reassigns labels to the cells based on objective measurements until the classifier converges to a stable grouping proposal.

To perform objective classification, the neuronal morphologies must be encoded in a digital format. The 3D digital reconstruction of a neuron encodes the path (in XYZ co-ordinates) and the thickness of each branch within its morphology and enables the consistent morphological analysis of its structure. The standard morphometrics (such as section length, bifurcation angles, etc.; Petilla Interneuron Nomenclature Group 2008, Ascoli & Krichmar 2000) that are commonly used as input measurements for objective classification, focus on different local aspects of the neuronal morphology and therefore must be used in combination with other morphological measurements. To avoid over-fitting, i.e., confusing the random noise in the biological structure with a significant discrimination factor, which is a result of using a large number of features in a few individual cells, feature selection is required. Appropriate feature selection is important for identifying the features that are indicative of the differences between neuronal shapes and that can be generalized across different brain regions and species. However, feature selection is often subjective, and the feature sets proposed by different experts are often inconsistent (DeFelipe et al. 2013). In addition, alternative sets of morphometrics result in different classifications (Kanari et al. 2017). To avoid this issue, a number of mathematically rigorous methods have been proposed for the morphological analysis of neurons (Van Pelt et al. 1991, DeFelipe et al. 2013, Gillette and Ascoli 2015a, Gillette et al. 2015B, Wan et al. 2015). We have developed an alternative representation of morphologies based on persistent homology (Carlsson, 2009) that provides a standardized quantification of neuronal branching structure (Kanari et al. 2017).

The Topological Morphology Descriptor (TMD) algorithm generates a barcode from a neuronal tree, coupling the topology of the branching structure with its geometry, and therefore encoding the overall shape of the tree in a single descriptor (Kanari et al. 2017). The TMD is a simplified representation of the original tree that retains key information to perform well in a discrimination task, by mapping the tree to a topological representation with less information loss than the usual morphometrics. The cell types proposed based on the TMD-classification are unbiased, since they are based on a mathematical descriptor of the tree’s branching structure rather that the visual inspection of the cells, and thus this method is less prone to user-induced biases. There is no need to analyze the tree based on different morphological features in an attempt to use and combine the ones that are significant. This way we avoid over-fitting by implicitly accounting for the correlations between features that are incorporated into their TMD profile.

Using this topological representation, we were able to establish that PCs can be objectively classified based on the branching structure of their apical dendrites. We compared the results of the topological classification to the expert-proposed cell types and illustrate that the majority of subjective cell types can be objectively supported, with the exception of L5 subtypes (TPC_A and TPC_B) and the rare horizontal PCs that are found in L6.

## Materials and Methods

### Staining and reconstruction techniques

All animal procedures were approved by the Veterinary Authorities and the Cantonal Commission for Animal Experimentation of the Canton of Vaud, according to the Swiss animal protection law.

The 3D reconstructions of biocytin-stained PC morphologies were obtained from whole-cell patch-clamp experiments on 300 μm thick brain slices from juvenile rat somatosensory cortex, following experimental and post-processing procedures as previously described (Markram et al. 1997b). The neurons that were chosen for 3D reconstruction were high contrast, completely stained, and had few cut arbors. Reconstruction used the Neurolucida system (MicroBrightField Inc., USA) and a bright-field light microscope (DM-6B, Olympus, Germany) at a magnification of 100x (oil immersion objective, 1.4-0.7 NA) or of 60x (water immersion objective, 0.9 NA). The finest line traced at the 100x magnification with the Neurolucida program was 0.15 μm. The slice shrinkage due to staining procedure was approximately 25% in thickness (Z-axis). Only the shrinkage of thickness was corrected at the time of reconstruction. Reconstruction resulted in a set of connected points traced from the image stacks of the 3D neuronal morphology, each having a 3D (X, Y, Z) position and diameter. The reconstructed PCs from all layers of rat somatosensory cortex were then used for the topological classification.

### Visualization of morphologies

The reconstructed morphology skeletons are represented by connected sets of points that account for the neurites and two-dimensional profiles of somata. Accurate visualization of these morphologies requires simulating the three-dimensional profile of the soma. We used a recent method to build a surface mesh model of the entire skeleton that represents its surface membrane. This method simulates the soma growth using Hooke’s law and mass-spring systems (Abdellah et al., 2017).

### Topological classification

For the topological classification, we first separated the PCs into layers according to the location of their somata, defined by somata positions as labeled by experts during the reconstruction process. The PCs were then separated into cell types based on the topological morphology descriptor (TMD) of their apical branching patterns (Kanari et al. 2017). The same analysis shows no significant differences among the branching patterns of basal dendrites of different m-types. The local axons cannot be used for the classification as they are usually cut, and large parts of them are not represented accurately in the neuronal reconstruction. The branching pattern of each apical tree is decomposed into a persistence barcode (Carlsson 2009), which represents each branch of the tree as a pair of points that correspond to a selected morphological feature.

The feature that is used as a filtration function influences the result of the classification. It is therefore important to select the morphological feature that will serve as the filtration function intelligently— based on the objective of the classification scheme—or to combine the TMD profiles for different features. In this study, we used the radial distance from the soma as the discriminating factor (see also Kanari et al. 2017). The use of alternative features revealed that the use of the radial distance as a filtration function performs equivalently well as or better than a number of other features (such as path distance, section lengths etc.) and therefore there is no need to combine multiple functions for the current study. Because we want to take into account the orientation of the trees, the radial distance is weighted according to the orientation of the tree towards the pia surface. The algorithm and the details of the method that transforms the neuronal trees into the persistence barcodes were previously described in detail in Kanari et al. 2017. The TMD of the tree is then used for the generation of the *persistence image* (Chepushtanova et al. 2015) of the tree, which summarizes the density of components at different radial distances from the soma. The persistence image representation is a vector that can be used as input to various machine learning algorithms.

We first performed a supervised classification using the cell types assigned by experts. An *objective supervised classifier* is trained on the cell types proposed by experts. Each neuron is then labeled according to its TMD profile. The accuracy of the classification is the total number of TMD-labels that agree with the initial label over the total number of cells. The classification is then repeated for a set of randomized labels corresponding to the initial number of cell types. If the expert classification accuracy is significantly higher than that of the randomized classification, the proposed grouping is accepted. If this is not the case, the classification cannot be confirmed by the TMD. The cell types are then redefined according to the TMD profiles of the neurons of the same layer, with the objective of an optimal separation between the defined cell types.

### Reclassification

The objective of the TMD-based classification is to test whether the expert types present distinct branching patterns and to explain the differences between them. However, in some cases the TMD-classification on the expert-proposed cell types indicates the existence of a large number of misclassified cells. This mismatch is caused either by the fact that these types differ in features orthogonal to the branching of the neurons, which are not captured by the TMD, or by the fact that the expert classification, which is prone to human error, does not take into account the branching. In the second case, the grouping can be improved by a semi-supervised classification. Starting from the initial cell types, the misclassified cells are re-evaluated and distributed in new groups until the new grouping is stable. The reclassification is valid if the automatically assigned groups are better separated than the expert groups. If an optimal grouping is available, an objective clustering scheme can be proposed to replace the manually assigned types.

A measurement for the assessment of misclassification is the *confusion matrix*. In the context of supervised classification, the *confusion matrix*, also known as *error matrix* (Stehman, 1997) is used to illustrate the performance of an algorithm. The name “confusion matrix” stems from the fact that it makes it easy to see if the system is confusing two types (i.e., commonly mislabeling one as another). Each row of the matrix represents the predicted type while each column represents the expert type (or vice versa). The value of the matrix at row I, column J is the percentage of cells that are of type J, according to the experts, while the classifier predicts that they are of type I. If all predicted labels are the same as the expert labels, the diagonal of the matrix will be 1 (100% accuracy), and the rest of the matrix will be 0, indicating that the total error is 0%.

### Accuracy of classification

The accuracy of the TMD classifier reported in this study is based on the distance between the *persistence images* (Chepushtanova et al. 2015) of the apical dendrites of the neurons. However, when reclassification is required, the accuracy cannot be computed in terms of the same distance, to avoid over-fitting. In this case, the accuracy of the TMD-based classification is evaluated by the computation of a number of differents topological distances between a pair of persistence diagrams derived from the neuronal trees. These distances are not entirely independent from the distance defined between persistence images, but they capture properties of the diagrams that have not been taken into account in the TMD-classifier. Therefore, the evaluation of the TMD-classifier based on a number of “independently” computed distances will be more impartial. To evaluate TMD-classification performance, we computed the *bottleneck distance* (Edelsbrunner and Harer 2008), the *Wasserstein distance* (Villani 2003, Edelsbrunner 2010), the *sliced-Wasserstein distance* (Carriere et al. 2017), and a number of distances on the vectorized persistence diagrams: the distance between *landscapes* (Bubenik 2015) and the distance between *signatures* (Carriere et al. 2015). These distances are described in detail in the SI. The reported accuracy is computed as the average accuracy of the classification based on these distances.

## Results

We have characterized five major types of PCs, in agreement with expert observations. BPC (bitufted pyramidal cells, found in L6) are identified by two apical trees extending to opposite directions: one towards the pia and one towards the white matter. IPC (inverted pyramidal cells, found in L2 and L6) are identified by the main direction of their apical tree, which is oriented towards the white matter. HPC (horizontal pyramidal cells, found in L6) are also identified by the direction of their apical tree, which is oriented parallel to the pia, as opposed to all other types. The rest of the pyramidal cell types are oriented towards the pia: TPC (tufted pyramidal cells, found across all layers), which are identified by a distinct tuft formation, distal from the soma; UPC (untufted pyramidal cells, found in deep layers 4, 5, 6), which lack a clear tuft formation but extend to large radial distances; and SSC (spiny stellate cells, found in L4), which also lack a tuft formation, but also extend to small radial distances. In addition, three subtypes of TPC cells have been identified (A, B and C). TPC_A cells are the largest tufted cells and form a distinct large tuft (highest density of branches) and multiple obliques. TPC_B cells form a proximal tuft (close to the soma) that is larger than the tuft of TPC_C cells but have few or no obliques. TPC_C (previously termed slender-tufted pyramidal cells, Markram et al. 2015) form a small distal tuft and multiple obliques. The results of the topological analysis are described in the following section, organized by layers.

**Figure 1.**
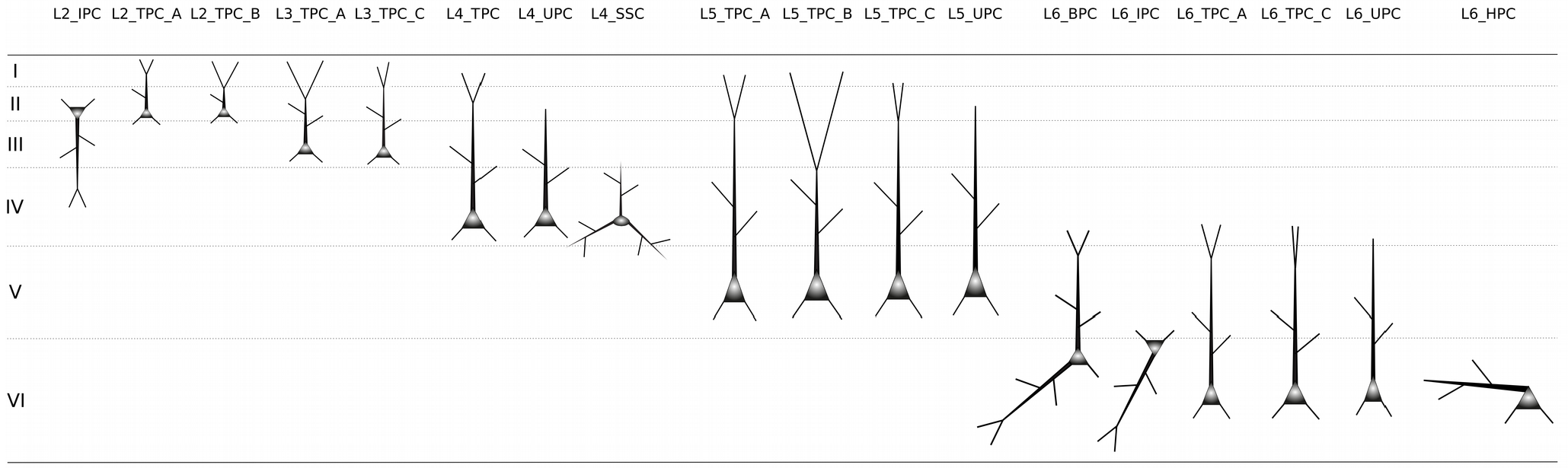
Schematic representation of all PC types/subtypes in Layers 2-6. Layer 2 consists of two main types of PCs, the L2_IPC (inverted PCs) and the L2_TPC (tufted PCs) which can be divided into two subtypes (_A – large tufted PCs, and _B – early bifurcating PCs). Layer 3 has one major type L3_TPC and two subtypes of PCs (_A – large tufted PCs, and _C – small tufted PCs). Layer 4 PCs are grouped into three types (L4_TPC; tufted PCs, L4_UPC; untufted PCs and L4_SSC; spiny stellate cells). Layer 5 PCs consist of two major types (L5_TPC; tufted PCs and L5_UPC; untufted PCs) and two subtypes of L5_TPC (_A – large tufted PCs, also known as thick tufted PCs and _C – small tufted PCs, also known as slender tufted). Layer 6 consists of five major PC types: L6_BPC (bitufted PCs), L6_IPC (inverted PCs), L6_TPC (tufted PCs), L6_UPC (untufted PCs) and L6_HPC (horizontal PCs). Also, two subtypes of L6_TPC are identified (_A – large tufted PCs, and _C – small tufted PCs, also known as narrow tufted).

### PCs in Layer 2

The TMD clustering of L2 PCs (n=43, Figure 2) based on their apical trees illustrates the existence of three subtypes with accuracy 84% (this result is cross-validated with five additional topological distances which yield an average accuracy of 83%). The **L2_IPC**s (inverted PCs, n=4), which are directed towards white matter, have apical trees that project in the direction opposite to the pia, therefore generating a higher density of branches in this direction (Figure 2). On the contrary, the **L2_TPC**s (tufted PCs, n=39) contain apical dendrites that project towards the pia, therefore exhibiting a higher density of branches in this direction. Further analysis of the branching patterns of L2_TPCs results in a separation into two subtypes (**L2_TPC_A**, n=6 and **L2_TPC_B**, n=33) depending on the density of branches on the distal apical dendrites: L2_TPC_A have a small density of branches within the tuft, while L2_TPC_B do not.

A quantitative analysis based on the morphometrics of 3D reconstructions of the three subtypes of PCs (L2_IPC, L2_TPC_A, L2_TPC_B) revealed a small but not significant quantitative difference of average soma sizes; the average soma surface area of L2_TPC_B is larger (+10%) than the surface areas of L2_TPC_A and L2_IPC somata. The basal dendrites of L2 PCs subtypes share similar morphological features, and therefore no morphological difference can be quantitatively justified. The average total length, surface area and volume of the apical dendrites of L2_TPC_B cells are larger than those of L2_TPC_A and L2_IPC cells, reflecting their broader extents. In addition, L2_TPC_B axons extend further, resulting in larger total lengths and surface areas, suggesting the formation of dense local axonal clusters. However, the results about the axonal morphometrics are inconclusive due to the significant loss of axonal mass described in the previous sections.

Expert-based observations of the same dataset suggest the existence of three distinct types. L2_IPC cells (inverted PC) have a vertically inverted apical dendrite projecting towards deep layers and white matter that forms a proximal or distal extensive tuft formation and multiple oblique dendrites. The apical dendrites of both L2_TPC_A and L2_TPC_B subtypes reach the pia and differ mainly in the bifurcating point along the apical dendrite where the tufts begin to form: proximal or distal. Therefore, the TMD-based classification supports the subjective observations for L2 PCs, and for consistency we use the expert-proposed terminology for those cell types.

**Figure 2.**
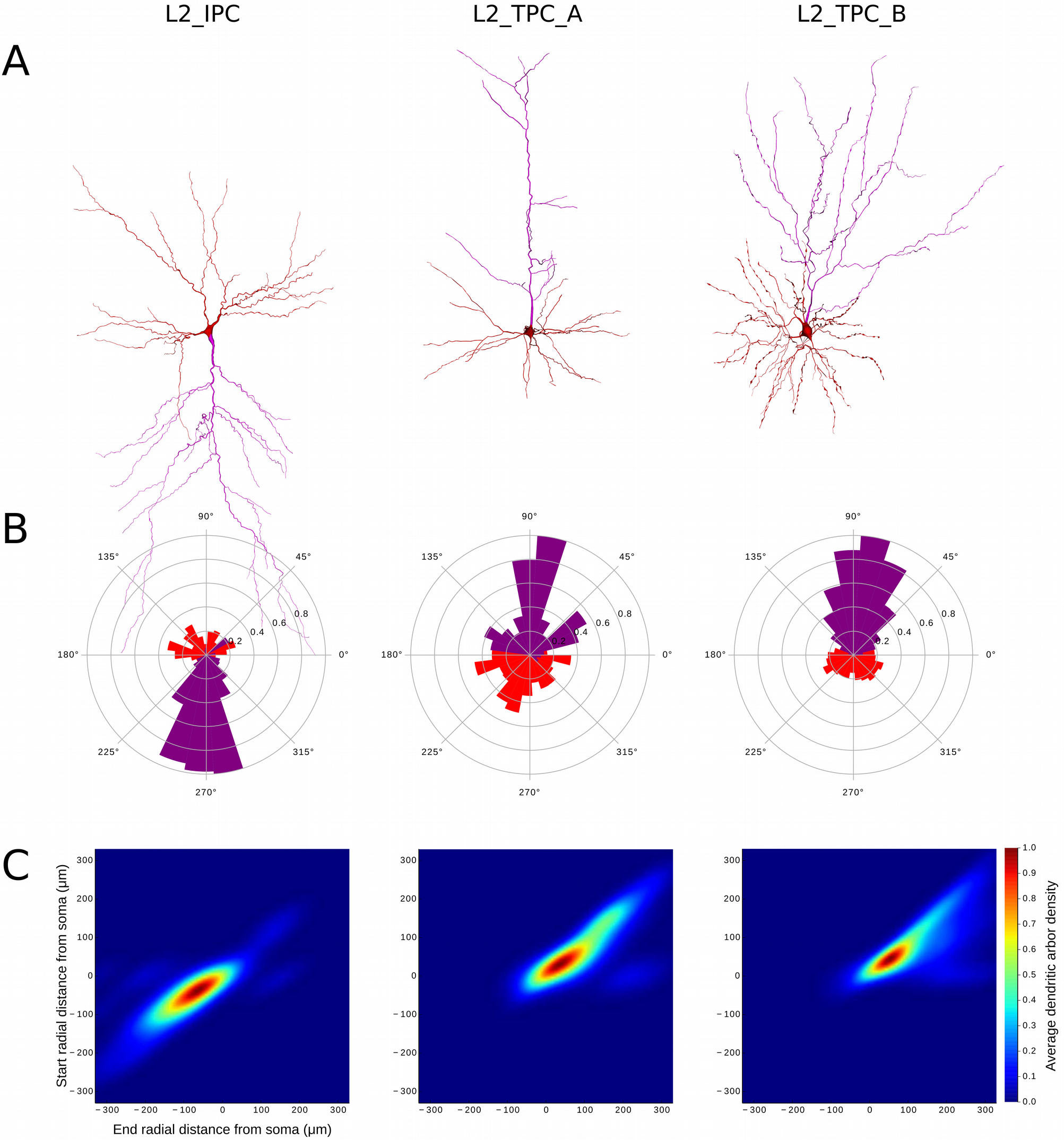
Three PC types/subtypes in Layer 2. A. Exemplar reconstructed morphologies of PC dendrites: the apical dendrite is presented in purple and the basal dendrites in red. B. Polar plot analysis of dendritic branches (apical in purple, basal in red). Tufted PCs are oriented towards the pia and the inverted PCs in the opposite direction as they project towards the white matter. C. The Topological Morphology Descriptor (TMD) of apical dendrites characterizes the spatial distribution of branches with respect to the radial distance from the neuronal soma. The average persistence images (per type of PC) illustrate the average dendritic arbor density around the soma. The spatial distribution of L2_IPC apical branches follow the direction opposite to the pia, while L2_TPC_A, which head towards the pia, present a larger density of branches for larger radial distances compared to L2_TPC_B.

### PCs in Layer 3

The TMD clustering of L3 PCs (n=44, Figure 3) based on their apical trees illustrates the existence of two subtypes with accuracy 89% (this result is cross-validated with five additional topological distances that yield an average accuracy of 85%). The **L3_TPC_A** (n=34) have apical trees with high density of branches close to the soma, but lower density of branches within the tuft. On the contrary, apical dendrites of **L3_TPC_C** (n=10) have a smaller density of branches around the soma, but higher density of branches on the tuft.

Quantitative morphological analysis on the two subtypes of L3 PCs (L3_TPC_A, L3_TPC_C) does not reveal any significant differences in the somatic and axonal features of the two subtypes. The differences between the two subtypes are captured only by the morphometrics of the apical dendrites. On average, L3_TPC_A cells have a larger number of oblique dendrites than L3_TPC_C cells, which corresponds to the lower densities of the latter observed in their persistence images. In addition, L3_TPC_A apicals have larger average lengths, surface areas, and volumes than L3_TPC_C.

Expert-based observations of the same dataset suggest the existence of two distinct types of L3 PCs, both of which are oriented towards the pia: the L3_TPC_A have a vertically projecting apical dendrite, with an often distal (occasionally proximal) onset of tuft formation, which forms a small tuft (occasionally extensive) and multiple oblique dendrites before tuft formation. On the contrary, the L3_TPC_C have a vertically projecting apical dendrite with distal onset of tuft formation, which forms a small tuft and few oblique dendrites before formation of the distal tuft. Therefore, the TMD-based classification supports the subjective observations of two subtypes in L3 PCs.

Compared to PCs in superficial layers (L2; Figure 2), L3 PCs appear to be larger on average, presenting larger extents and higher densities of branches, associated with larger total lengths. However, individual cells of L3 can be smaller than L2 cells, indicating that they cannot be distinguished merely by standard morphometrics. As a result, the information of soma location is essential for the analysis of L2 and L3 PCs. The axonal bouton density of L2 and L3 PCs is similar: with an average of 18–21 boutons/100 μm. Previous studies examined L2 and L3 PCs together, yielding two subtypes, which primarily differ in axonal morphology (Larsen and Callaway 2006) and therefore cannot be objectively linked to the cell types defined in this study, which are separated based on their apical dendrites. A subtype of superficial L2/3 PC sends axonal collaterals into L3 and 5, lacking axonal arbors in L4 while the other subtype, which is usually located at the bottom border of L3 (close to L4), has significantly more axonal collaterals within L4.

**Figure 3.**
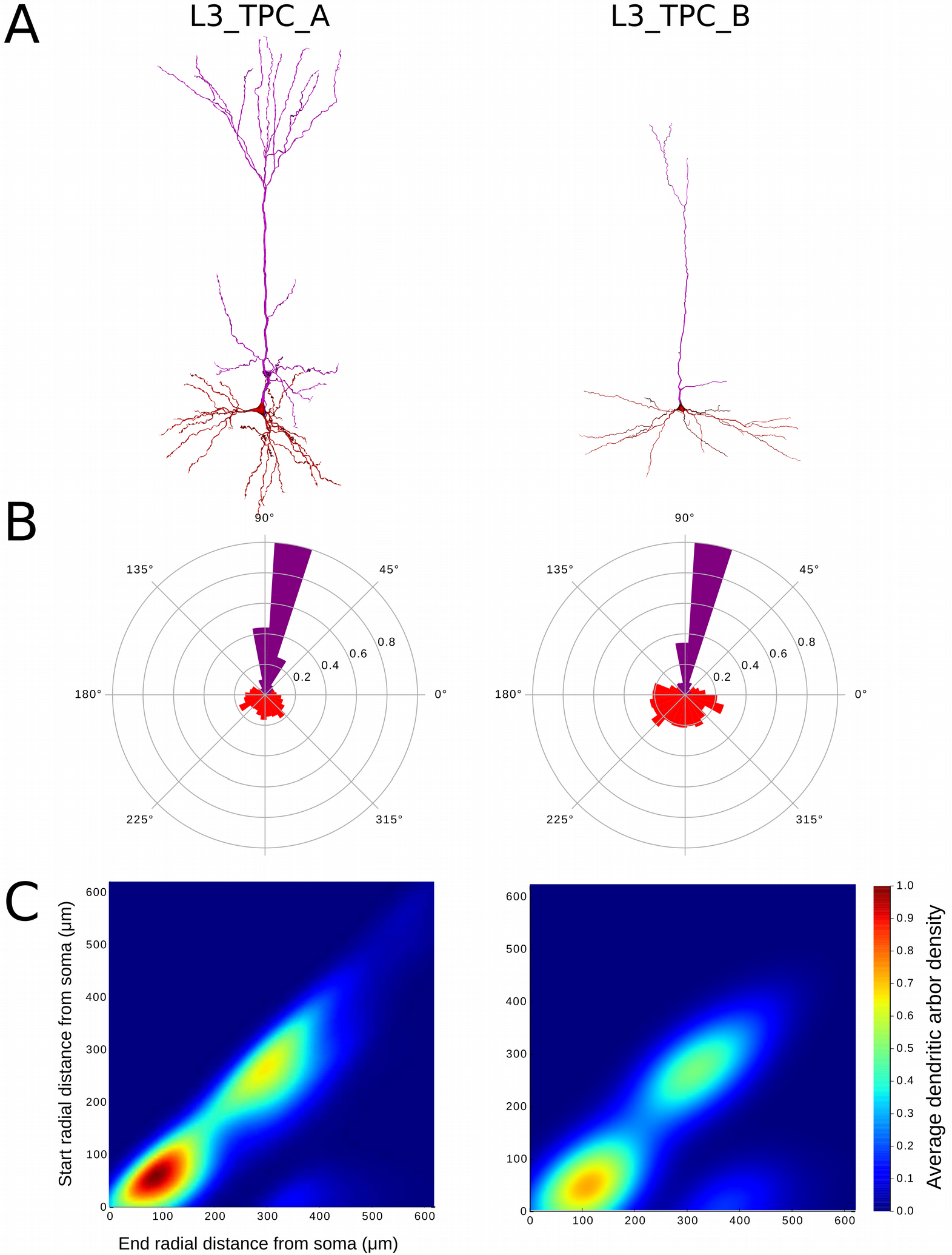
Two PC types/subtypes in Layer 3. A. Exemplar reconstructed morphologies B. Polar plot analysis of dendritic branches (apical in purple, basal in red). C. The Topological Morphology Descriptor (TMD) of apical dendrites characterizes the spatial distribution of branches with respect to the radial distance from the neuronal soma. The average persistence images (per type of PC) illustrate the average dendritic arbor density around the soma. L3_TPC_C apical dendrites present a lower density of branches both close to the soma and at larger radial distances. L3_TPC_A are denser and extend to larger radial distances.

The first case where reclassification is required is the L3 PC subtypes. Even though the score of the expert classification is high (86%), the confusion matrix (Figure 4C and 4D) indicates that the two subclasses are frequently confused for one another. In addition, a visual inspection of the cells (Figure 4A) illustrates that the two subclasses included cells with both large and smaller tufts and their structural differences are not easily identifiable.

**Figure 4:**
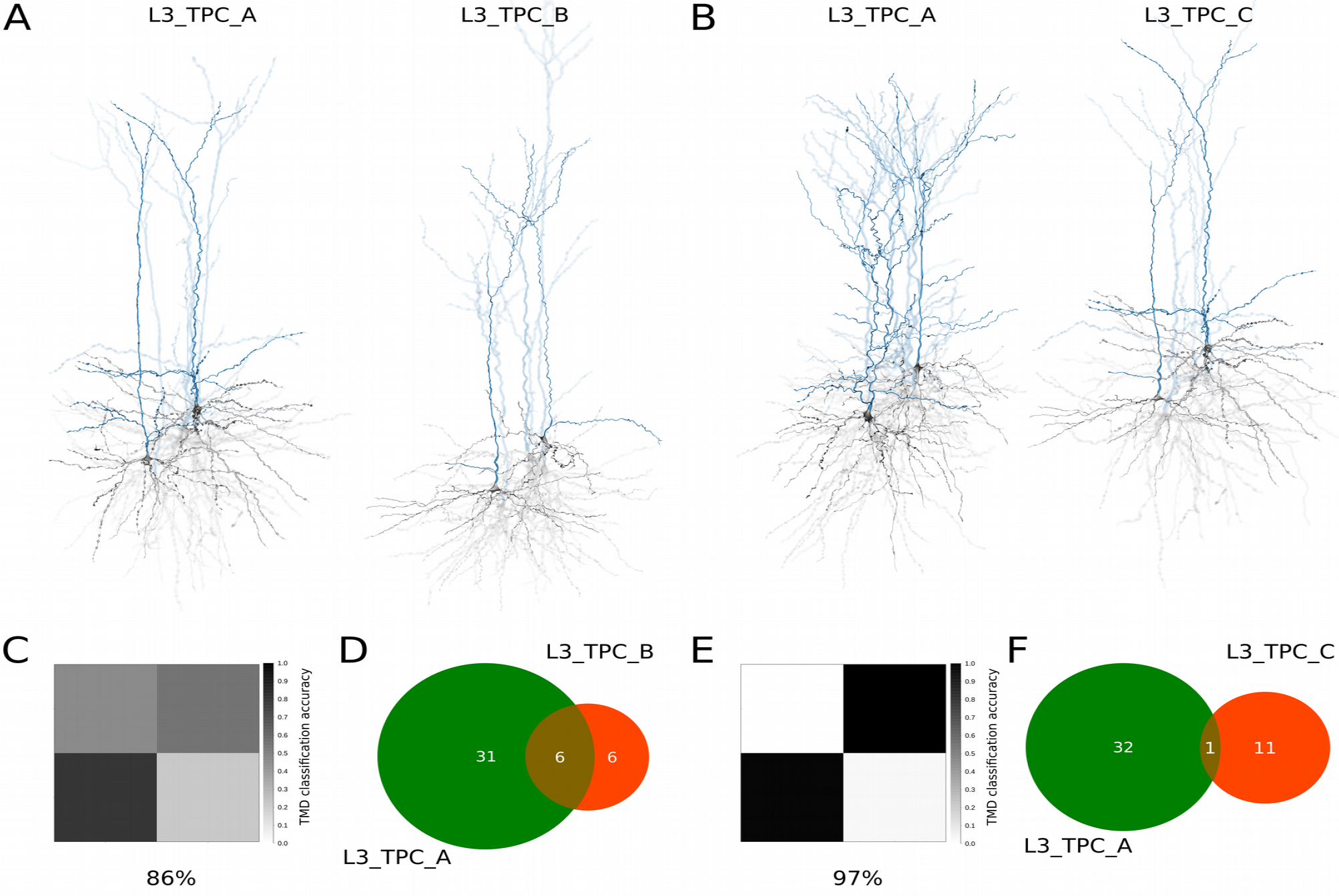
Reclassification of Layer 3 PCs. A. Curated renderings of L3_TPC_A and L3_TPC_B selected morphologies as proposed by expert classification. B. Curated renderings of L3_TPC_A and L3_TPC_B selected morphologies, after TMD-based reclassification. C. The confusion matrix illustrates the large percentage of misclassified cells between the expert proposed subtypes, yielding a total accuracy of 86%. D. The two subtypes are usually misclassified, as half of the L3_TPC_B are confused as L3_TPC_A. E. The confusion matrix illustrates the clear separation of the two subtypes after the TMD-based reclassification and the improved accuracy of the classifier (97%). F. The two subtypes are rarely misclassified, as almost all (~98%) of cells are unambiguously assigned into the two subtypes.

The reclassification according to the TMD profiles of their apical trees proposed a separation into two subtypes: the first includes cells with small tufted apical dendrites and the second cells with large tufted apical dendrites. This grouping is stable with respect to the automatic classification (Figure 4E and 4F), and visual inspection (Figure 4B) of the reclassified cells confirms that the two suggested subtypes correspond to L3_TPC_A (large tufted PCs) and L3_TPC_C (small tufted PCs). The expert grouping included a subtype (L3_TPC_B) in which half of the cells were unambiguous (Figure 4D). On the contrary the TMD-based clustering includes only one unambiguous cell (~2%, Figure 4F) yielding a well-defined separation of L3_TPC cells into the two proposed subtypes.

### PCs in Layer 4

The TMD clustering of L4 PCs (n=89, Figure 5) based on their apical trees illustrates the existence of two subtypes with accuracy 82% (this result is cross-validated with five additional topological distances that yield an average accuracy of 76%). The **L4_TPC**s (tufted PCs, n=44) have a long apical tree that extends to large radial distances and forms a tuft that presents a high density of branches at radial distances that are distal from the soma. The **L4_UPC**s (untufted PCs, n=33) apical trees also extend to large radial distances, but do not form a discrete tuft, as only few branches per tree reach the maximum radial distances. The apical trees of **L4_SSC**s (spiny stellate cells, n=12) present a high density of branches proximal to the soma, but only extend to small radial distances (about half of the radial distances of L4_TPCs).

Quantitative analysis based on the morphometrics of 3D reconstructions of the three subtypes of L4 PCs (L4_TPC, L4_UPC, L4_SSC) illustrates that L4_SSC have smaller somata than L4_TPC and L4_UPC. On average, compared to L4_UPC and L4_SSC, L4_TPCs have a larger number of basal dendrites, which are also significantly longer. Similarly, L4_TPCs apical trees are bigger (larger total length, areas and volumes) than both other types, even though both L4_TPC and L4_UPC apical extents are significantly longer than those of L4_SSC. Due to the significant loss of axonal mass, resulting from the slicing preparation (Stepanyants et al. 2009, Van Pelt et al. 2014), the results concerning the axonal morphometrics are inconclusive. However, the existence of three types is in agreement with previous studies that used thicker brain slices (500 μm thick) (Staiger et al. 2004) and reported three distinct types based on the axonal patterns of L4 PCs. In agreement with this study, the bouton density of L4_UPCs (22 ± 1 boutons/100 μm) is higher than those of L4_TPCs and L4_SSCs (19 ± 1 and 18 ± 1 boutons/100 μm respectively).

Expert-based observations of the same dataset suggest the existence of three types of L4 PCs, based on their apical dendrites. The L4_TPC (tufted PCs) have a vertically projecting apical dendrite with a small distal tuft and multiple oblique dendrites before tuft formation. The L4_UPC (untufted PCs) have a vertically projecting apical dendrite without a tuft and multiple oblique dendrites that branch proximally to the soma. The L4_SSC (spiny stellate cells) have a vertically projecting apical dendrite with small radial extents, not much longer than basal dendrites. Typically, the apical dendrites of all L4 PCs do not reach L1. Therefore, the TMD-based classification supports the subjective observations of three major types in L4 PCs.

**Figure 5.**
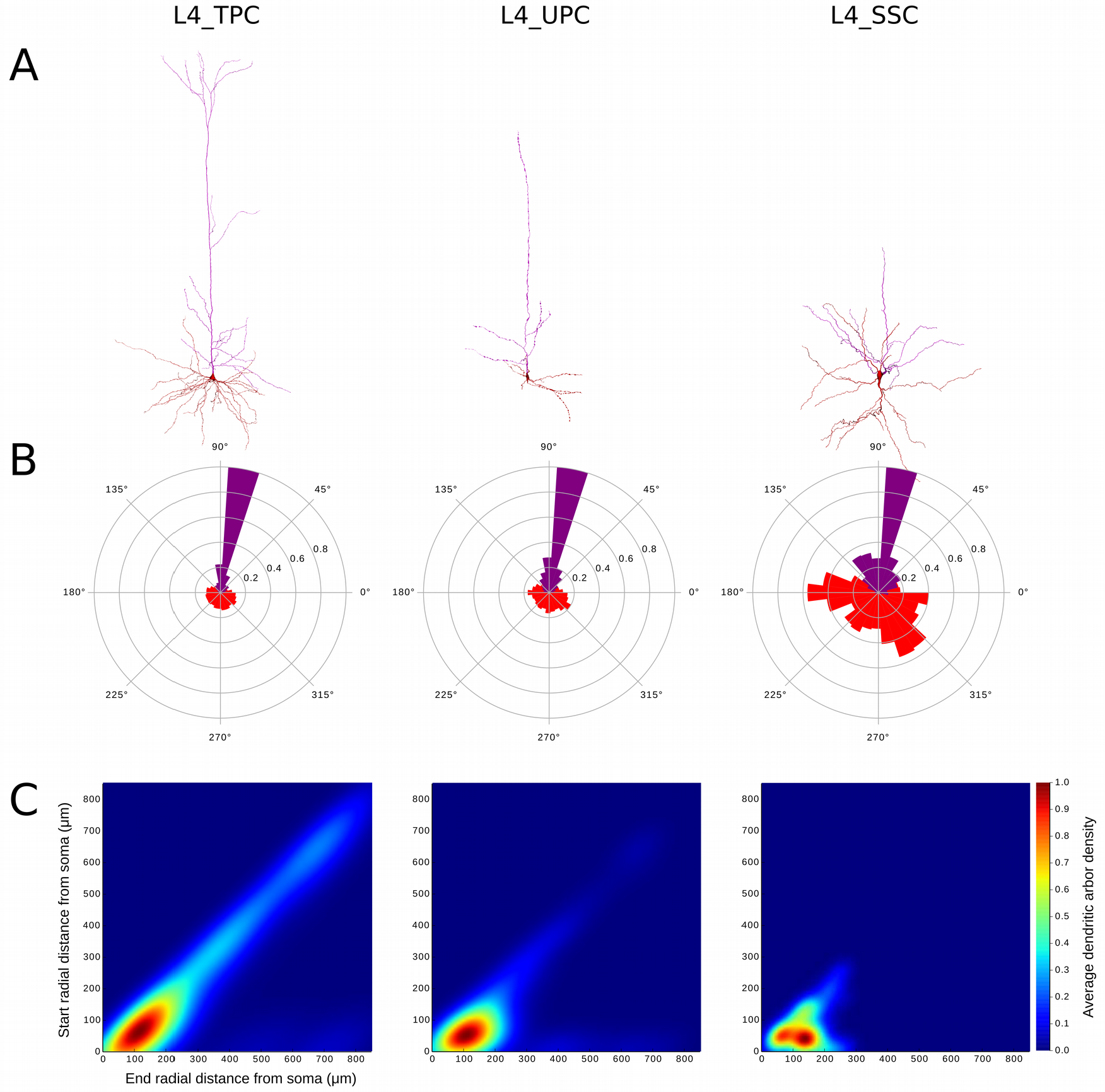
Three PC types/subtypes in Layer 4. A. Exemplar reconstructed morphologies B. Polar plot analysis of dendritic branches (apical in purple, basal in red). The polar plots of all types are similar as they all project towards the pia, with the exception of L4_SSC which remain local. C. The Topological Morphology Descriptor (TMD) of apical dendrites characterizes the spatial distribution of branches with respect to the radial distance from the neuronal soma. The average persistence images (per type of PC) illustrate the average dendritic arbor density around the soma. The apical dendrites of L4_TPC are the only ones that have a clearly defined tuft and are larger than both other types. L4_SSC are smaller than L4_UPC and their apical trees are similar to basal dendrites as they only extend to small radial distances.

### PCs in Layer 5

The TMD clustering of L5 PCs (n=160, Figure 6) based on their apical trees illustrates the existence of two subtypes with accuracy 90% (this result is cross-validated with five additional topological distances that yield an average accuracy of 84%). The TMD-based clustering of L5 PCs (n=160, Figure 6) based on their apical trees illustrates the existence of three subtypes of PCs that differ in the branching of their apical trees. The L5_TPCs (tufted PCs) can be objectively separated into two subtypes: A and C. **L5_TPC_A** cells (n=98) have a long apical tree that extends to the largest radial distances, reaching L1. L5_TPC_A apical trees have two distinct clusters with a high density of branches, which differ in their radial distance from the soma. The cluster proximal to the soma corresponds to the rich oblique formation, while the region distal from the soma corresponds to the formation of a densely branching tuft. Similarly, the apical dendrites of **L5_TPC_C** (n=32) have two distinct clusters of high branching density, one proximal to the soma that corresponds to the obliques and one distal to the soma that corresponds to the tuft. However, the tufts of L5_TPC_C have a lower density of branches, even though they extend to large radial distances. **L5_UPC** (untufted PCs, n=30) have a single high branching density cluster proximal to the soma, which corresponds to rich oblique formation. The reach of the apical trees of L5_UPC is lower than the rest of L5PCs, as the density of branches decreases with the radial distance from the soma, indicating the absence of a tuft.

**Figure 6.**
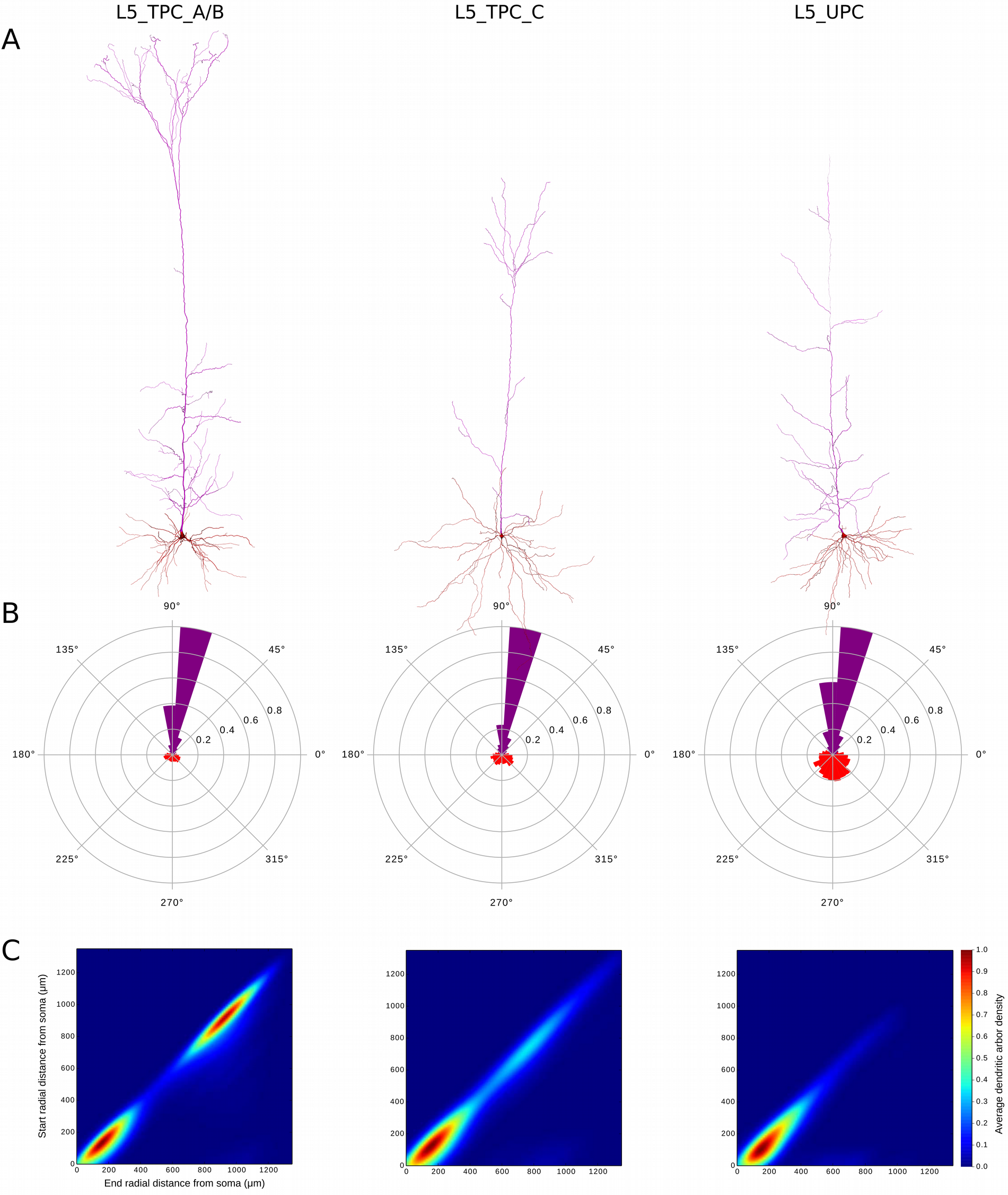
Three PC types/subtypes in Layer 5. A. Exemplar reconstructed morphologies B. Polar plot analysis of dendritic branches (apical in purple, basal in red). The polar plots of all types are similar as they all project towards the pia. C. The Topological Morphology Descriptor (TMD) of apical dendrites characterizes the spatial distribution of branches with respect to the radial distance from the neuronal soma. The average persistence images (per type of PC) illustrate the average dendritic arbor density around the soma. The apical dendrites of L5_TPC_A&B have a large tuft that extends to larger radial distances from the soma. L5_TPC_C similarly, present a tuft formation that is however smaller in size and extends to smaller radial distances. L5_UPC do not present a clear tuft but can also reach larger radial distances.

The quantitative analysis of 3D reconstructions of three subtypes of L5 PCs (L5_TPC_A, L5_TPC_C, L5_UPC) showed that L5_TPC_A have significantly larger somata compared to L5_TPC_C and L5_UPC. The basal dendrites of L5 PCs extend approximately to the width of a local cortical microcircuit (~300 – 500 μm), except those of L5_UPCs, which are narrower. L5_TPC_A have a significantly larger basal dendritic surface area, which enables higher synaptic inputs than the two subtypes that have longer but thicker basal processes. The morphological properties of L5_TPC_A apical trees confirm the topological results. In addition, L5_TPC_A cells (15-16 boutons/100 μm) have bouton densities significantly lower than those of L5_TPC_C and L5_UPCs (21 boutons/100 μm). Recent advances in retrograde labeling of single neurons *in vivo* with recombinant rabies virus (Larsen et al. 2007) resulted in the reconstruction of complete axons of L5 PCs, which supports the existence of three distinct subtypes based on their axonal properties. The thick-tufted PCs (corresponding to L5_TPC_A) project their local axons within deep layers, while the slender-tufted PCs (L5_TPC_C) and the short untufted PCs (L5_UPCs) have extensive projections to supra-granular layers. The axons of L5_UPCs are relatively columnar, while those of L5_TPC_Cs have extensive lateral spreads within L2/3. Compared to *in vivo* labeling (Larsen et al. 2007, Oberlaender et al. 2011), morphological measurements of axons obtained by *in vitro* (300 μm thick brain slices) labeling are underestimated, since the laterally spreading axonal processes are significantly severed during the slicing procedure.

Expert-based observations of the same dataset suggest the existence of two major cell types and four subtypes. The L5_TPC_A (thick-tufted PC_A) have a vertically projecting apical dendrite with a distal broad, thick tuft and multiple oblique dendrites emerging proximally. The **L5_TPC_B** (thick-tufted PC_B) are similar to the L5_TPC_A but further bifurcate into smaller tufts in comparison with L5_TPC_A. The L5_TPC_C (small-tufted PC) have a vertically projecting apical dendrite with a small distal tuft and multiple oblique dendrites emerging proximally. The L5_UPC (untufted PC) have a vertically projecting apical dendrite with no tuft formation.

The expert classification into types L5_TPC_A and L5_TPC_B could not be validated by the TMD-based clustering, as no significant differences were found in the topological profiles of those subtypes. A reclassification based on the topological profiles of their apical trees, showed that there is a gradient between those two subtypes, as defined by experts, rather than a clear separation into two distinct types (Figure 7). Carefully selected exemplars of the two L5_TPC subtypes show a clear divergence between them as their topological distance is significantly high (Figure 7, right). However, the topological distance between cells of the two subtypes gradually decreases (Figure 7, left) revealing a convergence between them. This analysis illustrates that the two subtypes belong to a continuum, rather than two distinctly separated types (Figure 7). Therefore, the TMD-based classification supports the existence of three major types of L5_PCs, but not their separation into L5_TPC_A and L5_TPC_B subtypes. Further information, complementary to their branching structure, is required for the distinction of those subtypes, but was not available at the time of this study.

**Figure 7.**
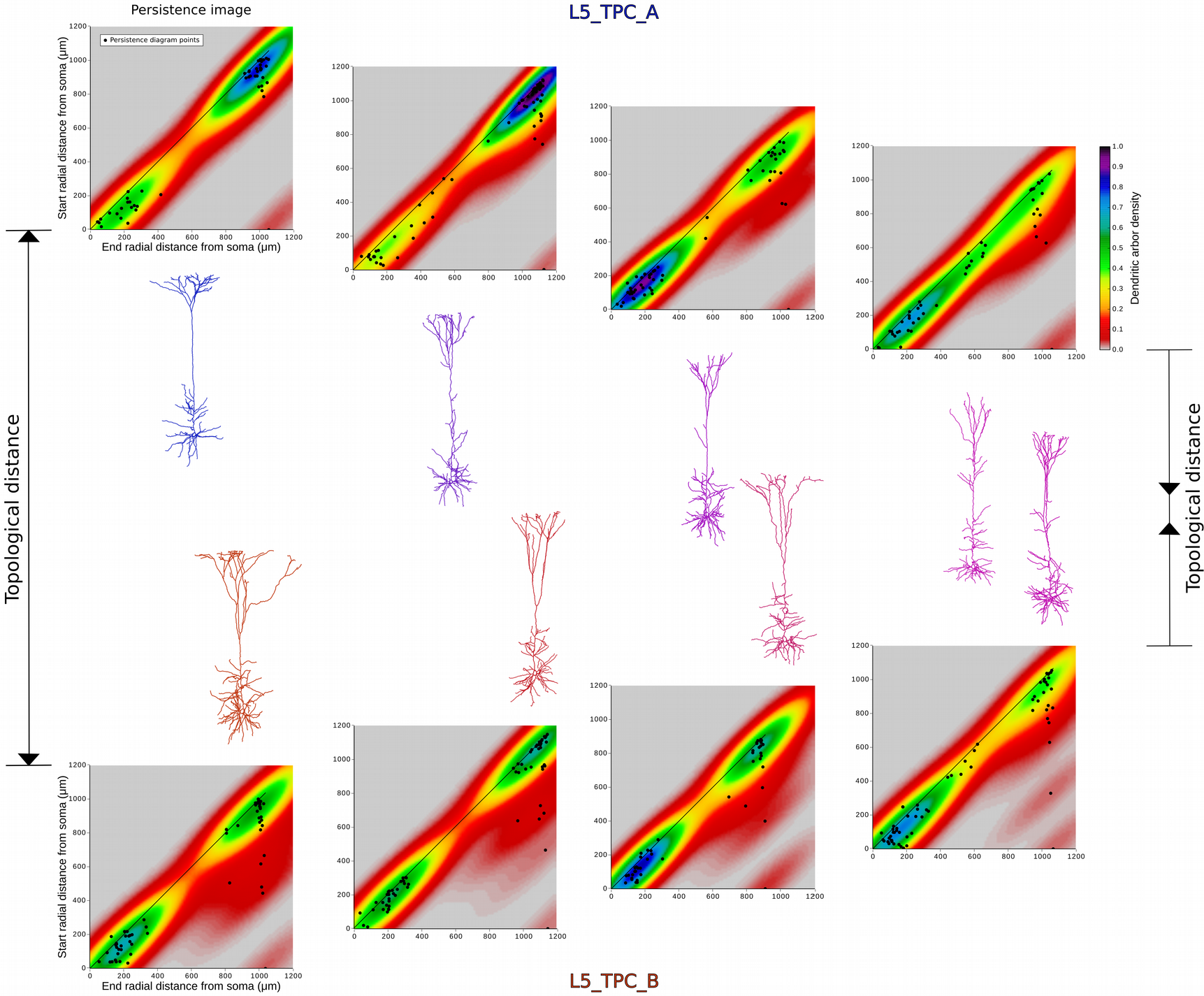
Convergence of subtypes L5TPC_A and B. Illustration of selected dendritic morphologies of L5_TPC_A (in blue) and L5_TPC_B (in red) of decreasing topological distance (from left to right). For border cases the two subtypes are very well separated (extreme left). The persistence images of all the presented apical trees are shown, and the points of the persistence diagrams for each apical tree are superimposed on the respective persistence images. However, as the topological distance decreases as the persistence images converge (left to right), and morphologies exhibit similar topological shapes (extreme right).

### PCs in Layer 6

The TMD clustering of L6 PCs (n=123, Figure 8) based on their apical trees illustrates the existence of two subtypes with accuracy 92% (this result is cross-validated with five additional topological distances that yield an average accuracy of 72%). **L6_BPC** cells (bitufted PCs, n=32) are identified by two vertically projecting branching clusters that project to opposite directions. Both of the apical trees of L6_BPC form a small distal tuft, which is indicated by a small distal cluster of branches in the persistence image (Figure 8), and a high density of branches close to the soma. **L6_IPC** (inverted PCs, n=26) are identified by the orientation of their apical trees, which are directed towards white matter. The low distal branching density of the L6_IPC apical indicates the existence of a small tuft. L6_TPC (tufted PCs, n=49), which form a distinct, large tuft, can be separated into two subtypes, as in the case of L5 PCs. The **L6_TPC_A** cells (n=22) have a long apical tree that extends to large radial distances (and reaches L4) and forms two clusters of branches at different radial distances from the soma. The cluster proximal to the soma corresponds to the rich oblique formation, while the distal cluster corresponds to the formation of a densely branching tuft. The **L6_TPC_C** cells (n=27) also have two distinct clusters of branches, one proximal to the soma that corresponds to the obliques and one distal to the soma that corresponds to the tuft. However, the tufts of L6_TPC_C have a lower density of branches than L6_TPC_A. **L6_UPC** (untufted PCs, n=16) apicals have a single dense cluster of branches proximal to the soma, which corresponds to a rich oblique formation. L6_UPC have smaller extents than L6_TPC, and the density of branches decreases with the radial distance from the soma, indicating the absence of a tuft. Note that even though the four proposed types (and two subtypes of L6_TPC) are consistent with the expert observations (see below), the individual cells were reclassified into these groups according to their TMD profiles because the expert classification was based only on visual observations. The reclassification redistributed the misclassified cells and confirmed the existence of the expert-proposed groups with similar properties. The last subtype of L6 PCs is **L6_HPC** (horizontal PCs). This subtype cannot be identified with the TMD-based classifier, as the apical dendrites of L6_HPC have similar topological profiles to the L6_UPCs. However, L6_HPC have a preferred horizontal orientation, as opposed to all the other L6 PCs, and therefore they can be objectively distinguished if the main direction of the apical tree is taken into account.

The quantitative analysis based on the morphometrics of 3D reconstructions of L6 PCs (L6_TPC_A, L6_TPC_C, L6_UPC, L6_IPC, L6_BPC, L6_HPC) shows that the somata of L6_HPCs are the biggest in L6 compared to other subtypes. L6_TPC_C basal dendrites are the smallest (minimum total length) among all L6 PCs, while the L6_HPCs basal dendrites have the widest maximum horizontal extent, but the smallest number of dendritic trees. L6_TPC_As and L6_UPCs have greater total dendritic length than all other L6 PCs, except HPCs. Quantitative analysis of L6 PCs axons demonstrates that they are largely similar, with the exception of L6_TPC_Cs, which have the narrowest axonal trees with the smallest maximum horizontal extent, which is approximately equal to the width of a cortical column. In addition, the L6_TPC_Cs have the lowest bouton density (17 boutons/100 μm) and the L6_HPCs the highest (22 boutons/100 μm). The other types/subtypes of L6 PCs all have similar bouton densities, ranging from 19 to 20 boutons/100 μm on average. Since a significant part of the axons of L6 PC reconstructions cannot be retrieved due to the slicing of the tissue, as discussed in previous sections, especially since L6 axons typically extend through multiple cortical columns (Boudewijns, Kleele et al. 2011), the morphometrics of L6 axonal branches will not be discussed further.

Subjective observations suggest the existence of five major types and two subtypes. The L6_TPC_A (tufted PC) have a vertically projecting apical dendrite with a small distal tuft and multiple oblique dendrites. The L6_TPC_C (narrow PC) have a narrow, vertically projecting apical dendrite, with a small distal tuft and often more oblique dendrites than other PC types. The L6_UPC (untufted PC) have a vertically projecting apical dendrite with no tuft formation, but multiple oblique dendrites. The L6_IPC (inverted PC) have a vertically inverted apical dendrite projecting towards the white matter with a small distal tuft and multiple oblique dendrites. The L6_BPC (bitufted PC) have two vertically projecting apical dendrites: one oriented toward the pia with a small distal tuft that forms multiple oblique dendrites and one inverted, projecting towards the white matter with a small distal tuft and multiple oblique dendrites. The L6_HPC (horizontal tufted PC) have a horizontally projecting apical dendrite with a small distal tuft that forms a few oblique dendrites. The apical dendrites of L6 PCs often reach L4 or supra-granular layers, but very rarely reach L1. Therefore, the TMD-based classification supports the existence of five subtypes in L6, and an additional cell type (L6_HPC) can be identified by using the main orientation of the apical tree as a distinctive parameter.

The types of L6 PCs identified in this study are in agreement with the proposed types of L6 PCs in a previous study (Marx and Feldmeyer 2012). L6_BPC correspond to the “multipolar neurons” of the study by Marx and Feldmeyer (2012), L6_IPC to “inverted neurons”, L6_TPC correspond to the “pyramidal cells”, L6_UPC to the “tangentially oriented neurons” and L6_HPC to the “horizontally oriented neurons”. Using this analogy as an example, we can identify potential links between the locally defined types of pyramidal cells and their respective long-range projections, as proposed in literature.

**Figure 8.**
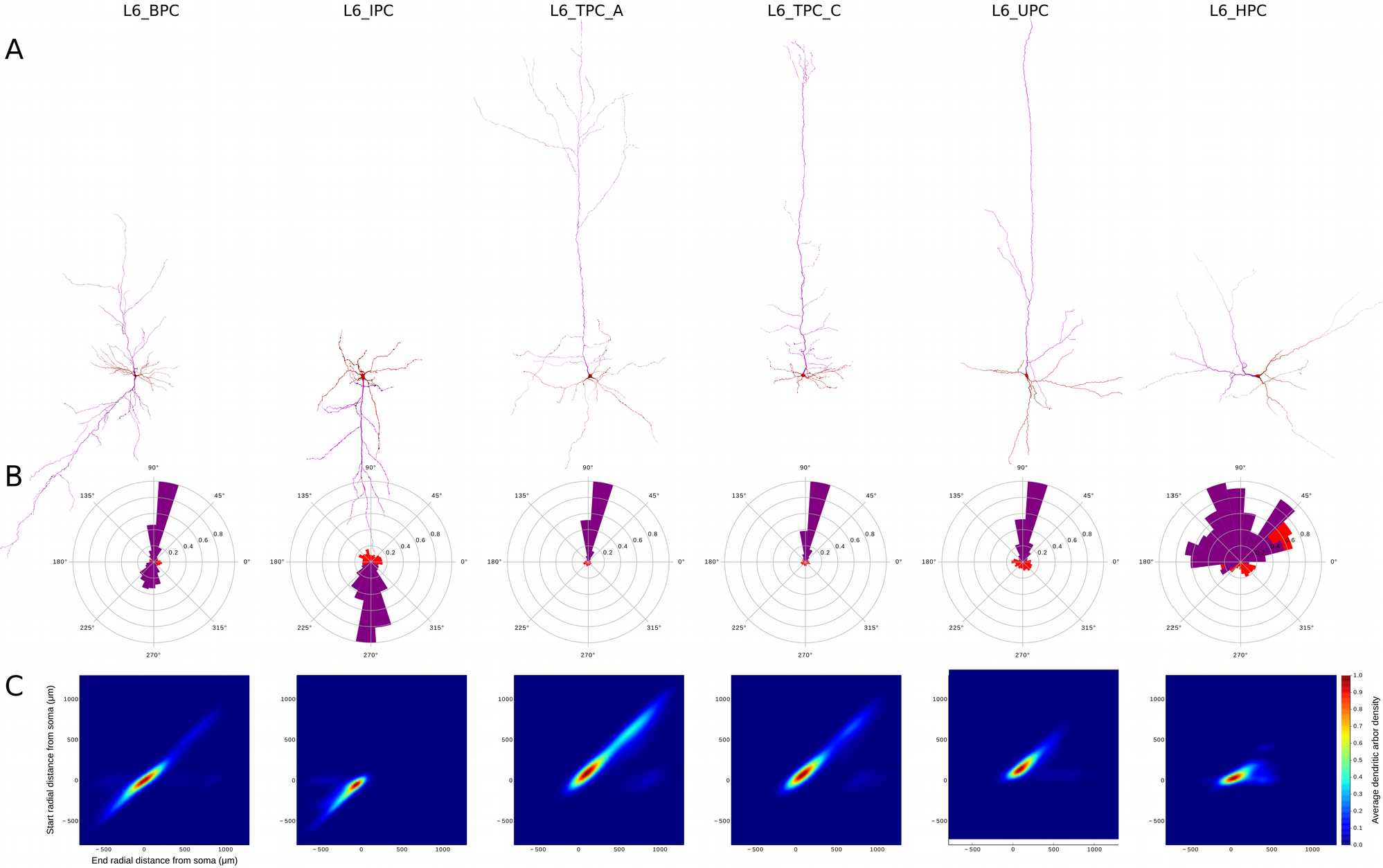
Six PC types/subtypes in Layer 6. A. Exemplar reconstructed morphologies of B. Polar plot analysis of dendritic branches (apical in purple, basal in red). The polar plots of L6_BPC and L6_IPC differ from other types that project towards the pia. Similarly, L6_HPC is the only type which presents almost symmetric polar plots due to their horizontal orientation. C. The Topological Morphology Descriptor (TMD) of apical dendrites characterizes the spatial distribution of branches with respect to the radial distance from the soma. The average persistence images (per type of PC) illustrate the average dendritic arbor density around the soma. L6_BPC have two distinct apical trees, that extend to opposite directions. Apical dendrites of L6_IPCs project away from the pia, towards the white matter. L6_TPC_As form a large tuft at large radial distances from the soma, while L6_TPC_Cs form a narrow tuft usually in smaller radial distances. Apical dendrites of L6_UPCs do not form a clear tuft but extend to large radial distances. L6_HPC is unique to layer 6 and projects vertically within the layer.

### Summary

The results of the TMD classification are summarized in Figure 9, where the percentage of occurrence of each m-type (Table 1, Figure 9A) and the corresponding accuracy of the classification (Figure 9B) are reported. The topological analysis of the branching structure of the PCs’ apical dendrites revealed the existence of sixteen subtypes of cells in all cortical layers, and one more subtype was objectively identified in L6 (Figure 10). The objective classification justifies the existence of three subtypes in L2 (Figure 2), two subtypes in L3 (Figure 3), three subtypes in L4 (Figure 5), three subtypes in L5 (Figure 6), and six subtypes in L6 (Figure 8). According to the laminar assignment of PCs by experts, no PCs were found in L1. The apical dendrites of PCs in supra-granular L2/3 reach L1 and the pia. The apical dendrites of PCs in layers 4 and 6 often reach the supra-granular layers, but not L1. Major PC subtypes in L5 have the longest apical dendrites, which reach L1 and the pia, and minor PC subtypes in L5 tend to extend to the supra-granular layers, but not to L1 (Figures 1 and 10).

**Figure 9:**
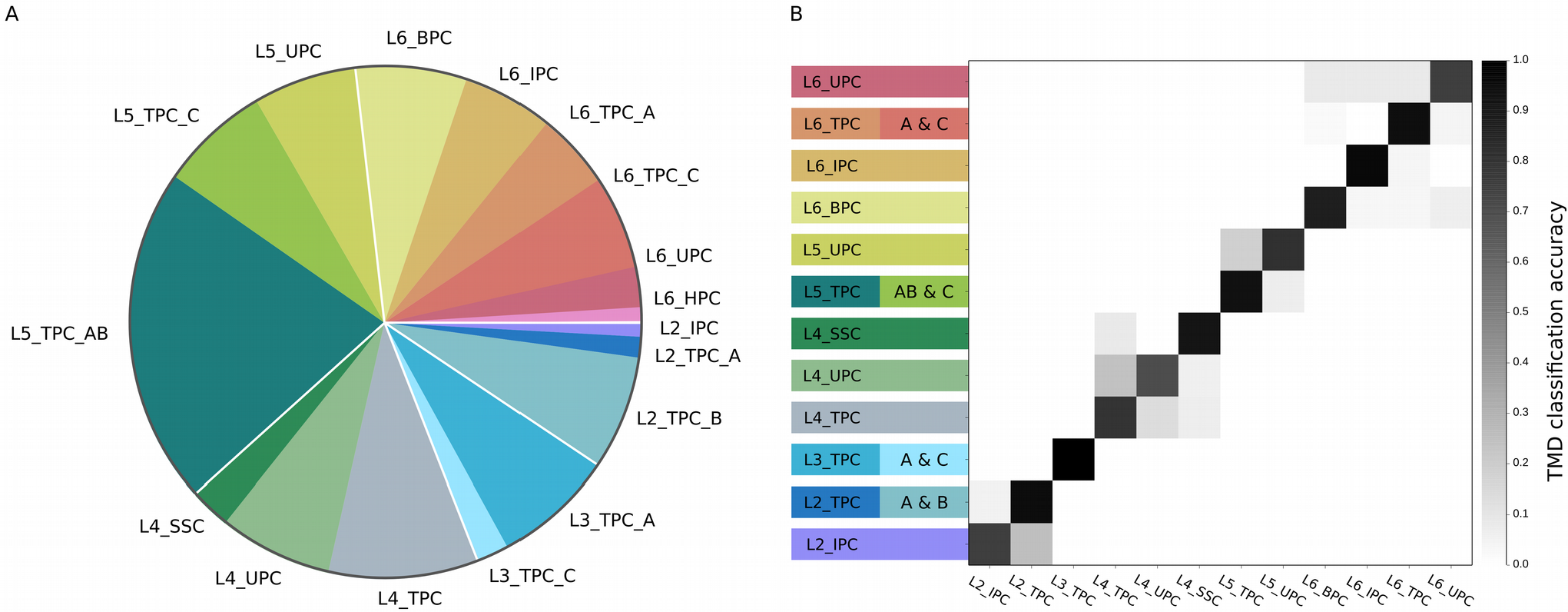
Summary of TMD-based classification of all cortical PCs. A. The pie-chart shows the percentage of cells per type/subtypes of cortical PCs, in different colors. The color-code is arbitrarily chosen. B. The confusion matrix illustrates the accuracy of the classification for the TMD-defined classes. The values on the diagonal show the percentage of cells for which the automatic and input labels agree and illustrate the accuracy of the classification (black – high accuracy; white – low accuracy). The perfect classification would have only black on the diagonal and white everywhere else.

The expert analysis of the cells revealed the existence of two subtypes that are not justified by the topological analysis: a subtype of TPC cells in L5, and a horizontally oriented cell type in L6. The L5_ TPC subtypes proposed by the experts are related to the visual characteristics of the cells but cannot be confirmed by our objective characterization of the cells. Instead of a rigorous separation between these two subtypes, the TMD reveals that there is rather a continuous gradient between them. The second type not supported by the TMD, the L6_HPC, can be distinguished by the main orientation of their apical dendrites, but the topological profiles of these cells are indistinguishable from the untufted L6 PCs (L6_UPC). A reclassification was required for the definition of subtypes in Layer 3 and Layer 5. In addition, L6_TPC_A, L6_TPC_C and L6_UPC were redefined according to the TMD classification.

**Figure 10:**
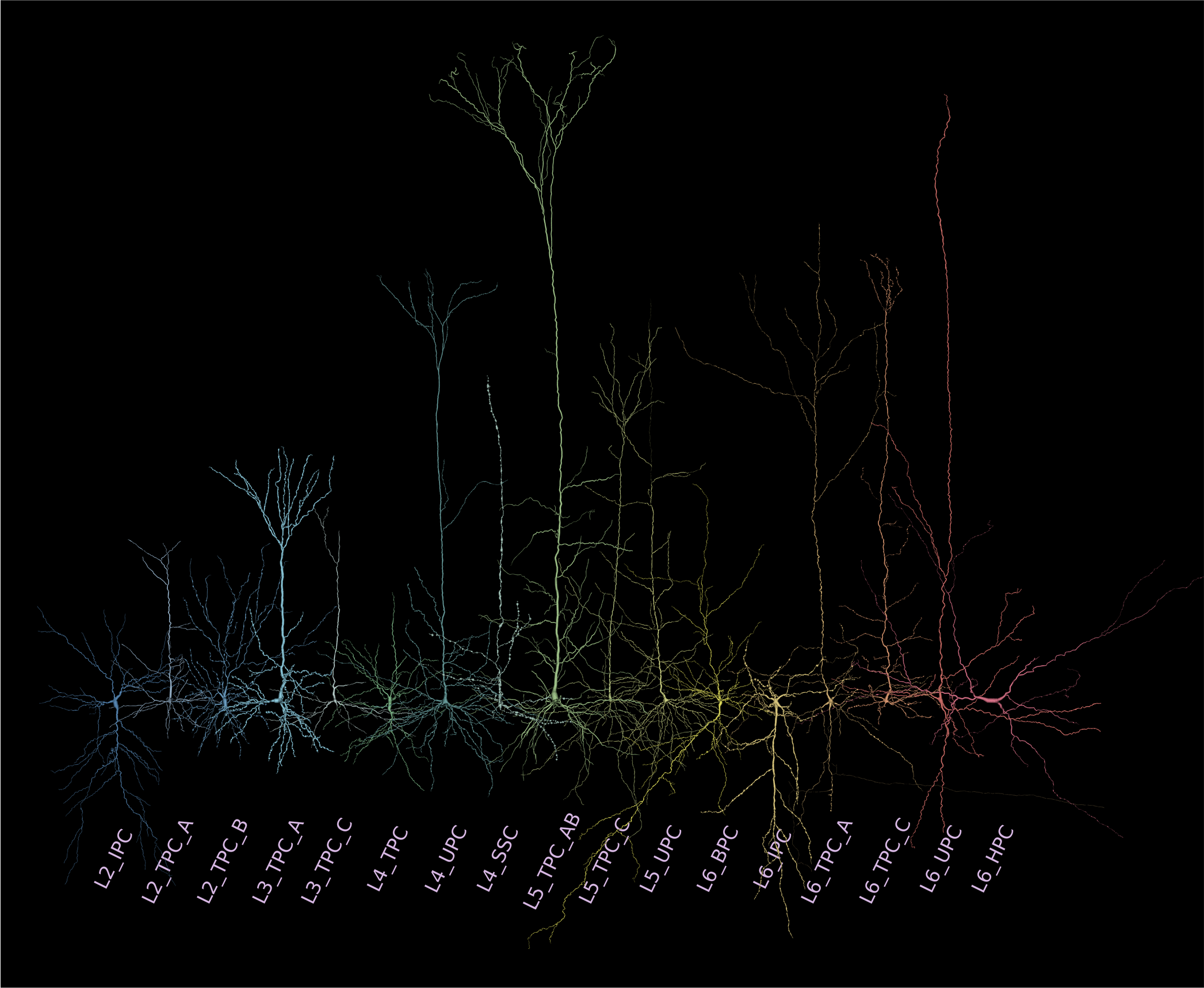
Overview of all cortical PCs types/subtypes. Renderings of dendrites and somata of all types/subtypes in order of appearance in the text. The color-code is arbitrarily chosen for consistency with Figure 9. Deeper layers express a larger diversity of PC types as the complexity of branching types increases from Layer 2 to Layer 6. The dendritic diameters have been scaled (x2) for better resolution of the dendritic morphologies.

**Table 1.** Number of PC types as identified by the TMD-based classification

| Layer | Type of PCs | Subtype of PCs | Number of PCs | Percentage |
| --- | --- | --- | --- | --- |
| 2 | IPC | - | 4 | 0.86 |
| 2 | TPC | A | 6 | 1.29 |
| 2 | TPC | B | 33 | 7.10 |
| 3 | TPC | A | 33 | 7.10 |
| 3 | TPC | C | 11 | 2.37 |
| 4 | SSC | - | 12 | 2.58 |
| 4 | TPC | - | 44 | 9.46 |
| 4 | UPC | - | 33 | 7.10 |
| 5 | TPC | A&B | 98 | 21.07 |
| 5 | TPC | C | 32 | 6.88 |
| 5 | UPC | - | 30 | 6.45 |
| 6 | BPC | - | 32 | 6.88 |
| 6 | HPC | - | 7 | 1.50 |
| 6 | IPC | - | 26 | 5.59 |
| 6 | TPC | A | 25 | 5.38 |
| 6 | TPC | C | 22 | 4.73 |
| 6 | UPC | - | 17 | 3.66 |

## Discussion

Despite the expertise involved, visual inspection of neurons is subjective and often results in non-consensual and ambiguous classifications (DeFelipe et al. 2013). In this study, we used a novel metric based on persistent homology (Kanari et al. 2017), which quantifies the branching structure of apical dendrites, to establish an objective standardized classification of pyramidal cells in the juvenile rat somatosensory cortex. We have demonstrated that the TMD of neuronal trees is not only reliable in validating the quality of the expert classification but can also propose an alternative separation of cells into groups when expert classification fails to provide a consensual and consistent definition of neuronal types.

Our classification scheme not only validates the existence of a common type of PCs across layers 2-6 (TPC) but also of several types that are unique to specific layers, such as the SSC in L4 and the BPC in L6. Interestingly, the diversity of shapes of apical dendrites increases with the distance from the pia, indicating that the higher functional complexity of deeper cortical layers can be successfully supported by the large morphological variability that is present in deeper layers, in agreement with recent observations (Reimann et al. 2017). The TMD-based classification was unable to distinguish a few cell types proposed by experts that differ in morphological characteristics that do not directly contribute to the branching structure, such as L6_HPC, which are distinguished by their horizontally oriented dendrites. In this particular case, an additional descriptor, i.e., the main orientation of the cell, was used for the objective discrimination of L6_HPC neurons. This demonstrates that expert classification is essential to guide further improvements of the method. The initial expert classification is also important for the training of the classifiers in order to automatically assess the type of a new cell based on a dataset of cells from previously validated groups.

Certain tools of the new subfield of algebraic topology called topological data analysis (TDA, Carlsson and Zomorodian, 2009) enable the study of multidimensional persistence of features and could be used for combining independent morphological measurements not currently considered in the computation of the TMD into multidimensional barcodes. Using this technique, independent characteristics could be combined into a single topological descriptor to strengthen even further its discriminative power. For example, cells that differ on parameters that are currently not considered, such as the thickness of the processes and the bouton density and cannot be distinguished with the TMD descriptor, could be discriminated by an extended multidimensional descriptor.

Another important characteristic that has not been included in this study and should ideally be combined in an improved version of the TMD descriptor is the projection patterns of axons, which are particularly important for long-range axons that target distant brain regions. A growing body of evidence suggests a strong correlation between locally defined types of PCs and their target regions, which are genetically determined early on during differentiation and prior to the migration of the neurons to their destination layers (Larkman and Mason 1990; O’Leary and Koester 1993, Kasper et al. 1994; Franceschetti et al. 1998; Gao and Zheng 2004; Larsen and Callaway 2006; Morishima and Kawaguchi 2006; Kumar and Ohana 2008; Marx and Feldmeyer 2012). Indeed, long-range axonal projection of PCs is an important feature that enables different computational functions and should therefore be taken into account for their classification (Larsen and Callaway 2006; Larsen et al. 2007; Hattox and Nelson, 2007; Brown & Hestrin, 2009; Boudewijns et al. 2011).

Due to technical limitations, the long-range projections of PCs are not currently available for a sufficiently large number of cells to allow for their systematic characterization. However, recent advances in optical imaging and long-range axon labeling techniques are enabling a systematic reconstruction of single neurons at the whole-brain level (Yuan et al. 2015, Gong et al. 2016). Hopefully, these advances will lead to the systematic characterization of whole-cell reconstructions, in order to quantify their long-range axonal projection properties and associate them to their local dendritic properties.

## Acknowledgements

This study was supported by funding from the ETH domain to the Blue Brain Project, the European Union Seventh Framework Programme (FP7/2007-2013) under grant agreements 604102 (HBP) and 720270 (HBP SGA1), and the National Natural Science Foundation of China (Grant No. 31070951). We thank Dr. Eilif Muller and Dr. Julian Shillcock for providing critical input to the study.

